# Dalpiciclib Partially Abrogates ER Signaling Activation Induced by Pyrotinib In HER2^+^HR^+^ Breast Cancer

**DOI:** 10.1101/2022.12.07.519433

**Authors:** Jiawen Bu, Yixiao Zhang, Nan Niu, Kewei Bi, Lisha Sun, Xinbo Qiao, Yimin Wang, Yinan Zhang, Xiaofan Jiang, Dan Wang, Qingtian Ma, Huajun Li, Caigang Liu

## Abstract

**Background:** Recent evidences from clinical trials (NCT04486911) revealed that the combination of pyrotinib, letrozole and dalpiciclib exerted optimistic therapeutic effect to treat HER2^+^HR^+^ breast cancer, however, the underlying molecular mechanism remained further investigation.

**Methods:** Through the drug sensitivity test, the drug combination efficacy of pyrotinib, tamoxifen and dalpiciclib to BT474 cells were tested. The underlying molecular mechanisms were investigated using immunofluorescence, western blot analysis, immunohistochemical staining and cell cycle analysis. Potential risk factor which may indicate the responsiveness to drug treatment in HER2^+^/HR^+^ breast cancer was selected out using RNA-sequence and tested using immunohistochemical staining and in vivo drug susceptibility test.

**Results:** We found that pyrotinib combined with dalpiciclib exerted better cytotoxic efficacy than pyrotinib combined with tamoxifen in BT474 cells. Degradation of HER2 could enhance ER nuclear transportation, activating ER signaling pathway in BT474 cells whereas dalpiciclib could partially abrogate this process. This may be the underlying mechanism by which combination of pyrotinib, tamoxifen and dalpiciclib exerted best cytotoxic effect. Furthermore, CALML5 was revealed to be a risk factor in the treatment of HER2^+^/HR^+^ breast cancer and the usage of dalpiciclib might overcome this.

**Conclusion:** Our study provided evidence that the usage of dalpiciclib in the treatment of HER2^+^/HR^+^ breast cancer could partially abrogate the estrogen signaling pathway activation caused by anti-HER2 therapy and revealed that CALML5 could serve as a risk factor in the treatment of HER2^+^/HR^+^ breast cancer.

**Funding:** This study was supported by the National Natural Science Foundation of China (#U20A20381, #81872159)

## Introduction

Human epidermal growth factor receptor 2-positive (HER2^+^) breast cancer is associated with an increased risk of disease recurrence and death (*Perou et al., 2000; Slamon et al., 1987; Tzahar et al., 1996*). HER2-overexpressing breast tumors have high heterogeneity, accounting partially for the co-expression of hormone receptors (HR) (*Loi et al., 2016*). Previous studies have demonstrated that extensive cross-talk exists between the HER2 signaling pathway and the estrogen receptor (ER) pathway (*Wang et al., 2011*). In addition, exposure to anti-HER2 therapy may reactivate the ER signaling pathway, which could lead to drug resistance (*Brandao et al., 2020*). Generally, however, HER2-positive patients are treated using the same algorithms, both in the early and advanced stages (*Moja et al., 2012*).

Increasing evidence has confirmed that the intrinsic differences between HER2^+^/HR^+^ and HER2^+^/HR^-^ patients should not be ignored (*Carey et al., 2016*). Clinical outcomes have demonstrated that HER2^+^/HR^+^ breast cancer patients have a lower chance of achieving a pathologically complete response than HER2^+^/HR^-^ patients, when treated with neoadjuvant chemotherapy plus anti-HER2 therapy (*Cameron et al., 2017; Cortazar et al., 2014*). Nevertheless, the addition of concomitant endocrine therapy to anti-HER2 therapy or chemotherapy did not show any advantages in clinical trials, such as the NSABP B-52 and ADAPT HER2^+^/HR^+^ studies (*Harbeck et al., 2017*; *Rimawi et al., 2017*). Recently, the synergistic effect of CDK4/6 (cyclin kinase 4/6) inhibitors and anti-HER2 drugs in HER2^+^ breast cancer has been reported. The combination of anti-HER2 drugs and CDK4/6 inhibitors showed strong synergistic effects and high efficacy in HER2^+^ breast cancer cells (*Goel et al., 2016; Zhang et al., 2019*). Besides, in the recent MUKDEN 01 clinical trial (NCT04486911), the combination use of pyrotinib (anti-HER2 drug), letrozole (endocrine drug) and dalpiciclib (CDK4/6 inhibitor) exerted optimal therapeutic effect in HER2^+^HR^+^ breast cancer patients and offered novel chemo-free neoadjuvant therapy for the treatment of HER2^+^HR^+^ breast cancer (*Niu et al., 2022*), yet the underlying mechanism remained to be investigated.

Herein, we investigated the underlying molecular mechanism how the combination of pyrotinib, letrzole and dalpiciclib achieved satisfactory therapeutic effect in MUKDEN 01 trial. We studied the combined effect of pyrotinib (anti-HER2 drug), tamoxifen (endocrine therapy), and dalpiciclib (CDK4/6 inhibitor) on the HER2^+^/HR^+^ breast cancer cell line BT474 to simulate the clinical therapy in MUKDEN 01 trial(*Niu et al., 2022*). We found that pyrotinib combined with dalpiciclib exerted better cytotoxic efficacy than pyrotinib combined with tamoxifen. Moreover, the combination use of pyrotinib, tamoxifen and dalpiciclib displayed the best cytotoxic effect both in vitro and in vivo. In addition, HER2-targeted therapy induced nuclear ER redistribution in HER2^+^/HR^+^ cells and the activation of ER signaling pathway, which could be partially abrogated by the addition of dalpiciclib. Furthermore, the expression of CALML5 could be a potential risk factor in the treatment of HER2^+^HR^+^ breast cancer and the introduction of dalpiciclib could partially abrogate the drug resistance of HER2+HR+ breast cancer caused by the high expression of CALML5. Our study provided potential molecular mechanisms why the combination of pyrotinib, letrozole and dalpiciclib could achieve satisfactory clinical response and found CALML5 as a potential risk factor in the treatment of HER2^+^HR^+^ breast cancer.

## Results

### Pyrotinib combined with dalpiciclib shows better cytotoxic efficacy than when combined with tamoxifen

To explore the effects of anti-HER2 drugs, tamoxifen, and dalpiciclib in HER2^+^/HR^+^ breast cancer, we first evaluated the cytotoxic activities of these three reagents in BT474 breast cancer cells. The results indicated that the IC50 doses for pyrotinib, trastuzumab, tamoxifen, and dalpiciclib were 10 nM, 170μg/ml, 5 μM, and 8 μM, respectively (Figure 1-figure supplement 1a). To further investigate whether these drugs could have a synergistic effect in BT474 cells, we assessed the efficacies of the combinations of pyrotinib and dalpiciclib, pyrotinib and tamoxifen, and tamoxifen and dalpiciclib on the inhibition of cell proliferation at different concentrations. We calculated the combination index for each combination using Compusyn software to determine if the antitumor effects were synergistic (*Chou and Talalay, 1984*). Synergistic effects were observed in the combination group of pyrotinib and dalpiciclib, as well as in the pyrotinib and tamoxifen groups; both with CI values of <1 at several concentrations (Figure 1a). However, in the combination group of tamoxifen and dalpiciclib, no synergistic effect was observed.

**Figure 1.**
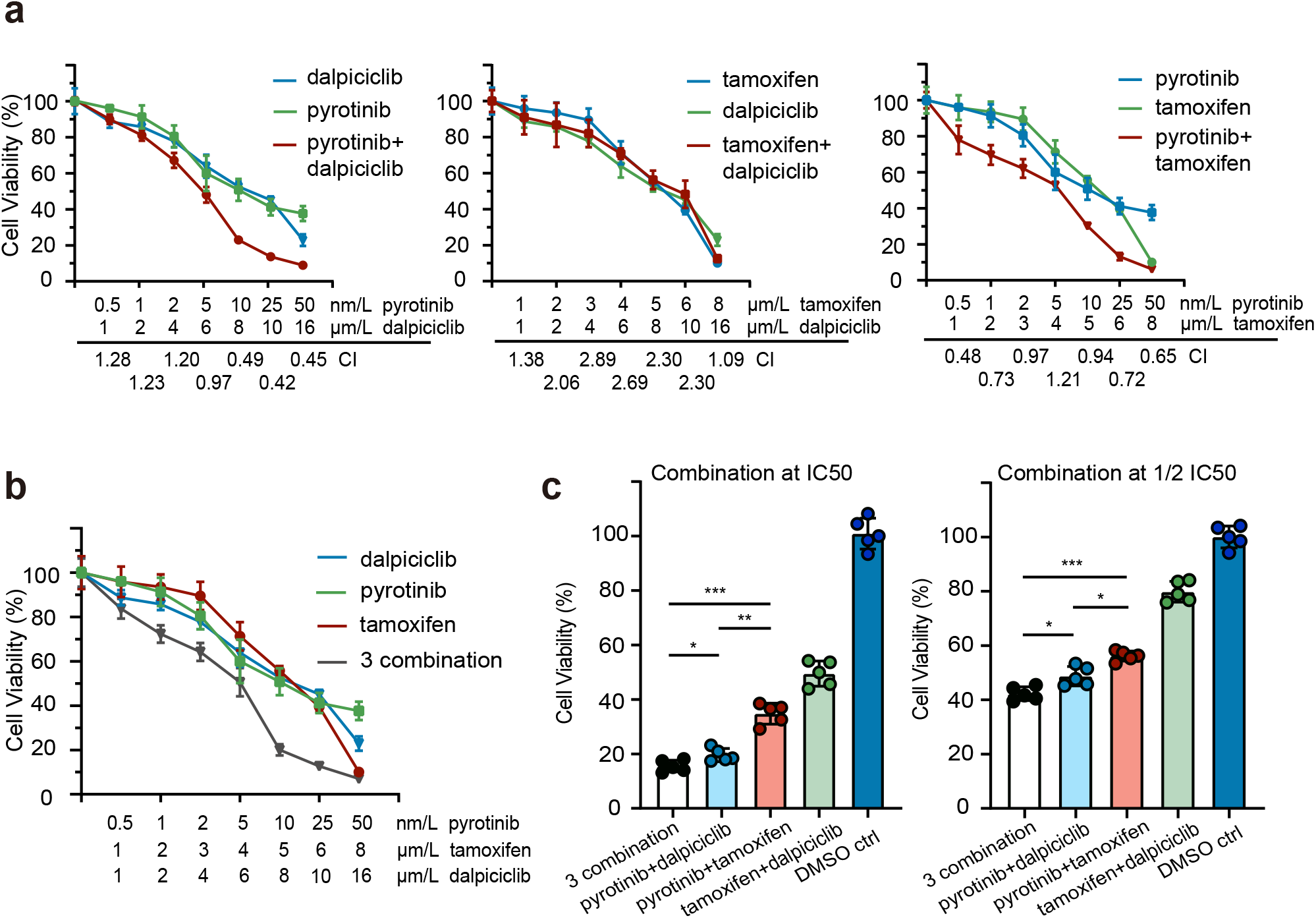
Drug sensitivity test of pyrotinib, tamoxifen, dalpiciclib and their combination on BT474 cells. a-b: Drug sensitivity assay of BT474 cells to single drug and different drug combination (Data was presented as mean ± SDs, all drug sensitivity assay were performed independently in triplicates). c: Drug sensitivity assay of BT474 cells to different drug combination at IC50 concentration and 1/2 IC50 concentration. (Data was presented as mean ± SDs, \**P*<0.05, \*\**P*<0.01 and \*\*\**P*<0.001 using repeated Anova test; all the assays were performed independently in triplicates) The statistical data was provided in Figure 1 source data 1.

We also analyzed the effect of the three-drug combination, and it showed a stronger cytotoxic effect on HER2^+^/HR^+^ breast cancer compared with the effect of the other two-drug combinations (Figure 1b). As both dalpiciclib and tamoxifen showed synergistic effects in combination with pyrotinib, we sought the combination that exerted better cytotoxic efficacy. Hence, we treated the BT474 cells with different combinations at IC50 or half IC50 concentrations. The three-drug combination and the combination of pyrotinib and dalpiciclib showed a stronger cell inhibition compared with that exerted by pyrotinib and tamoxifen as well as tamoxifen and dalpiciclib (Figure 1c). The colony formation assay also displayed similar trends as the cell viability assay; the three-drug combination formed the least number of colonies, followed by the combination of pyrotinib and dalpiciclib (Figure 1-figure supplement 1b).

### Nuclear ER distribution is increased after Anti-HER2 therapy and could be partially abrogated by the introduction of dalpiciclib

The results of the drug sensitivity test showed that the combination of pyrotinib and tamoxifen was less effective than the combination of pyrotinib and dalpiciclib. Considering that the expression of HER2 could affect the distribution of the ER(*Yang et al., 2004*), we performed immunofluorescence staining for ER distribution on the different drug-treated groups to see if anti-HER2 therapy could degrade HER2 and affect the distribution of ER. We found that pyrotinib induced ER nuclear translocation in BT474 cells, which could be partially abrogated by the addition of dalpiciclib, rather than tamoxifen (Figure 2a). Besides, trastuzumab, the monoclonal antibody of HER2 could also enhance the nuclear shift of ER and could also be abrogated by the introduction of dalpiciclib (Figure 2-figure supplement 1 c). Western blot analyses revealed although the total expression of ER was reduced, the nuclear ER levels increased considerably after the use of pyrotinib (Figure 2-figure supplement 1a–b). The use of tamoxifen increased the expression of total ER and nuclear ER (Figure 2-figure supplement 1a–b). However, when dalpiciclib was introduced, the increased expression of nuclear ER caused by pyrotinib was partially abrogated (Figure 2-figure supplement 1b) and this was consistent with the finding that dalpiciclib could increase the ubiquitination of ER (Figure 2-figure supplement 1d).

**Figure 2.**
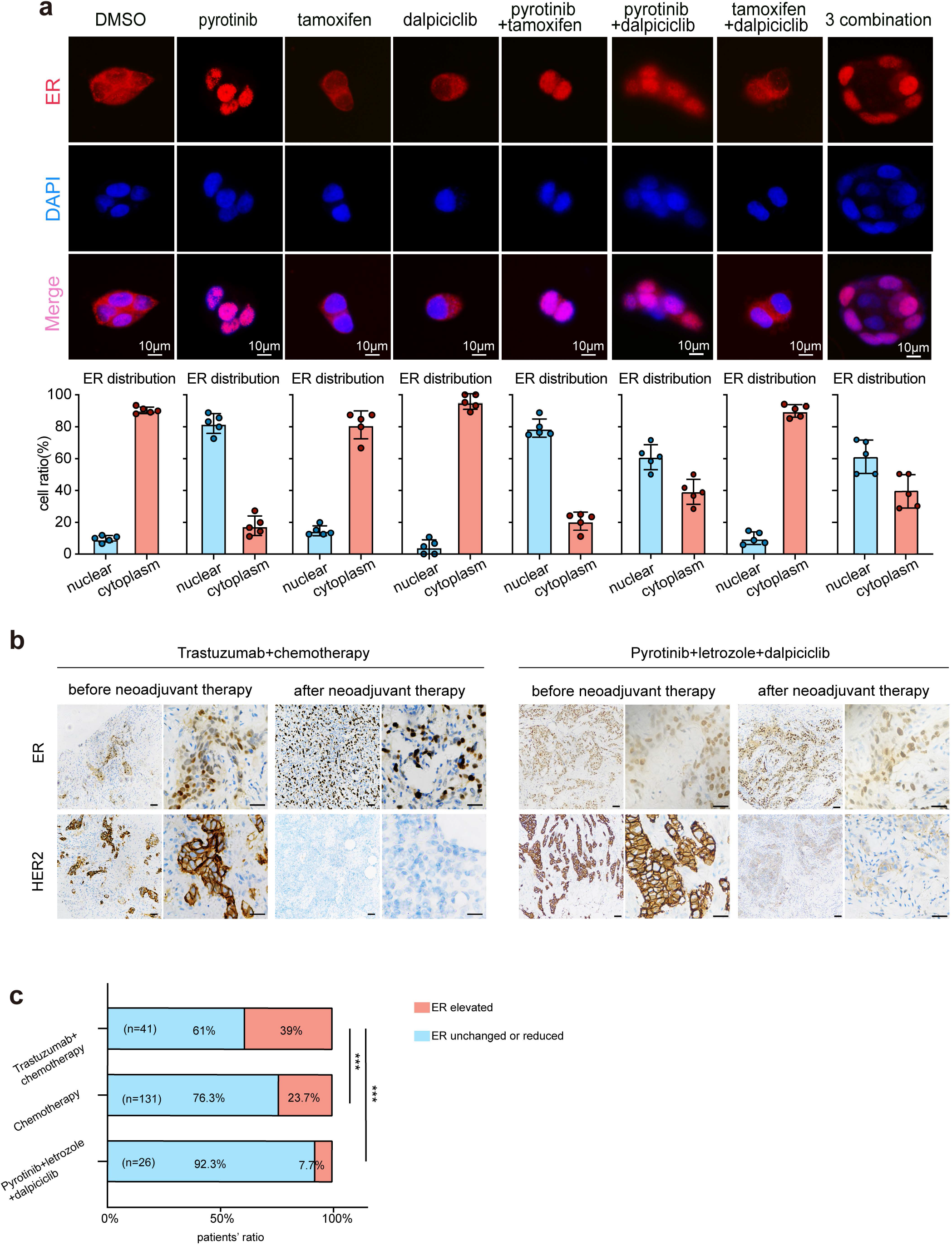
Anti-HER2 therapy could lead ER shifting into cell nucleus in HER2^+^/HR^+^ breast cancer while CDK4/6 inhibitor could reverse the nuclear translocation of ER. a: Distribution of estrogen receptor in BT474 cell line after different drug (pyrotinib, tamoxifen and dalpiciclib) treatment. (The distribution ratio of ER was calculated manually by randomly chosen 5 views in 400magnification. All the assays were performed independently in triplicates). b: Representative views of ER and HER2 expression in patients before and after anti-HER2 (trastuzumab) + chemotherapy (docetaxel+carboplatin) and representative views of ER and HER2 expression in patients before and after pyrotinib+letrozole+dalpiciclib treatment. c: Ratio of patients with elevated ER expression and patients with unchanged or reduced ER expression in different kinds of neoadjuvant therapy groups. (\*\*\**P*<0.001 using repeated Anova test) The statistical data was provided in Figure 2 source data 1.

Based on our in vitro findings, we further explored the ER distribution in clinical samples from the different treatment groups. To this end, we collected the clinical information of HER2^+^/HR^+^ patients who received neoadjuvant therapy at the Shengjing Hospital (Table 2). We found significant elevations in the nuclear ER expression levels of patients who received chemotherapy(doxetaxel+carboplatin) and anti-HER2 therapy (trastuzumab), compared with the levels in patients who only received chemotherapy (doxetaxel+carboplatin) (Figure 2b, c). However, in our clinical trial (NCT04486911, an open-label, multicentre phase II clinical study of pyrotinib maleate combined with CDK4/6 inhibitor and letrozole in neoadjuvant treatment of stage II-III triple positive breast cancer)(*Niu et al., 2022*), the nuclear ER expression levels of patients did not show significant elevations after the HER2-targeted therapy combined with dalpiciclib (Figure 2b, c). These findings verified that the ER receptor may have shifted to the nucleus after anti-HER2 therapy, which could be abrogated with the introduction of dalpiciclib.

**Table 1.**
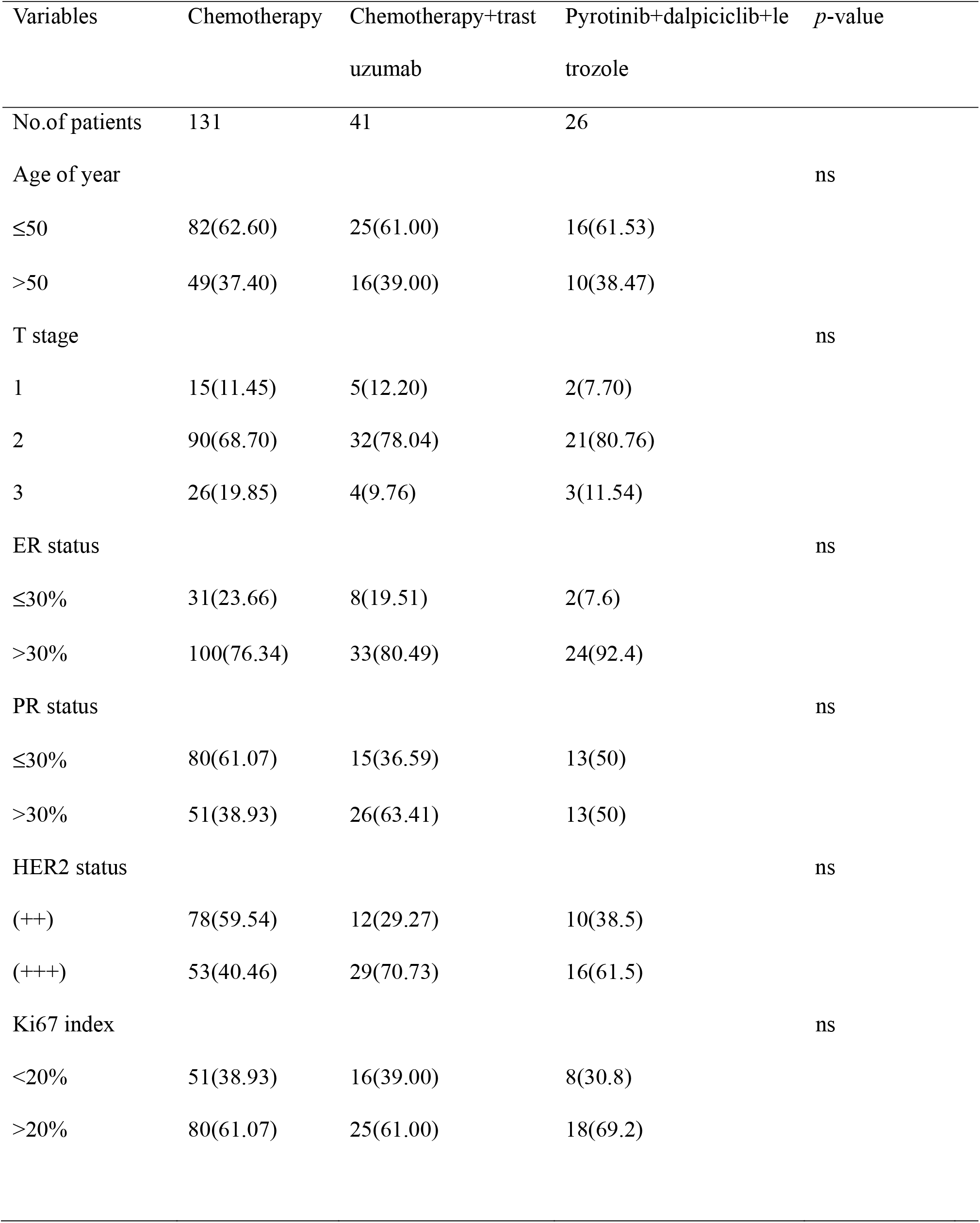
Demographic information of HER2^+^/HR^+^ breast cancer patients who received neoadjuvant therapy.

**Table 2.** Demographic information of HER2^+^/HR^+^ breast cancer patients who were tested for CALML5 before receiving neoadjuvant therapy.

| Variables | Chemotherapy+tras<br>tuzumab | Pyrotinib+dalpiciclib+let<br>rozole | *p*-value |
| --- | --- | --- | --- |
| No.of patients | 41 | 26 |  |
| Age of year |  |  |  |
| ≤50 | 25(61.00) | 16(61.53) | ns |
| >50 | 16(39.00) | 10(38.47) |  |
| T stage |  |  |  |
| 1 | 5(12.20) | 2(7.70) | ns |
| 2 | 32(78.04) | 21(80.76) |  |
| 3 | 4(9.76) | 3(11.54) |  |
| ER status |  |  |  |
| ≤30% | 8(19.51) | 2(7.6) | 0.0145 |
| >30% | 33(80.49) | 24(92.4) |  |
| PR status |  |  |  |
| ≤30% | 15(36.59) | 13(50) | ns |
| >30% | 26(63.41) | 13(50) |  |
| HER2 status |  |  |  |
| (++) | 12(29.27) | 10(38.5) | ns |
| (+++) | 29(70.73) | 16(61.5) |  |
| Ki67 index |  |  |  |
| <20% | 16(39.00) | 8(30.8) | ns |
| >20% | 25(61.00) | 18(69.2) |  |
| CALML5 |  |  |  |
| positive | 18(43.90) | 10(38.46) | ns |
| negative | 23(56.10) | 16(43.9) |  |

ns, not significant.

### Bioinformatic analyses unravel the synergistic mechanisms underlying the dalpiciclib and pyrotinib in HER2^+^/HR^+^ breast cancer

To further explore the mechanisms how dalpiciclib could partially abrogate the activation of ER signaling pathway after pyrotinib treatment, we first analyzed the gene expression profiles of the breast tumor cells treated with pyrotinib via RNA-seq. The signaling pathway enrichment analysis of the differentially expressed genes (DEGs) showed that majority of the DEGs were significantly enriched in the TNF signaling pathway and cell cycle, while steroid biosynthesis was also strongly active, suggesting that the steroid hormone pathway was activated by pyrotinib (Figure 3a-b). Similar results were obtained from the Gene Set Enrichment Analysis (GSEA). The administration of pyrotinib resulted in downregulation of the cell cycle and activation of the hormone pathway. The leading-edge subset of these pathways included the MITOTIC SPINDLE, G2M CHECKPOINT, and ESTROGEN RESPONSE EARLY (Figure 3c). These results showed good concordance with our in vitro findings.

**Figure 3.**
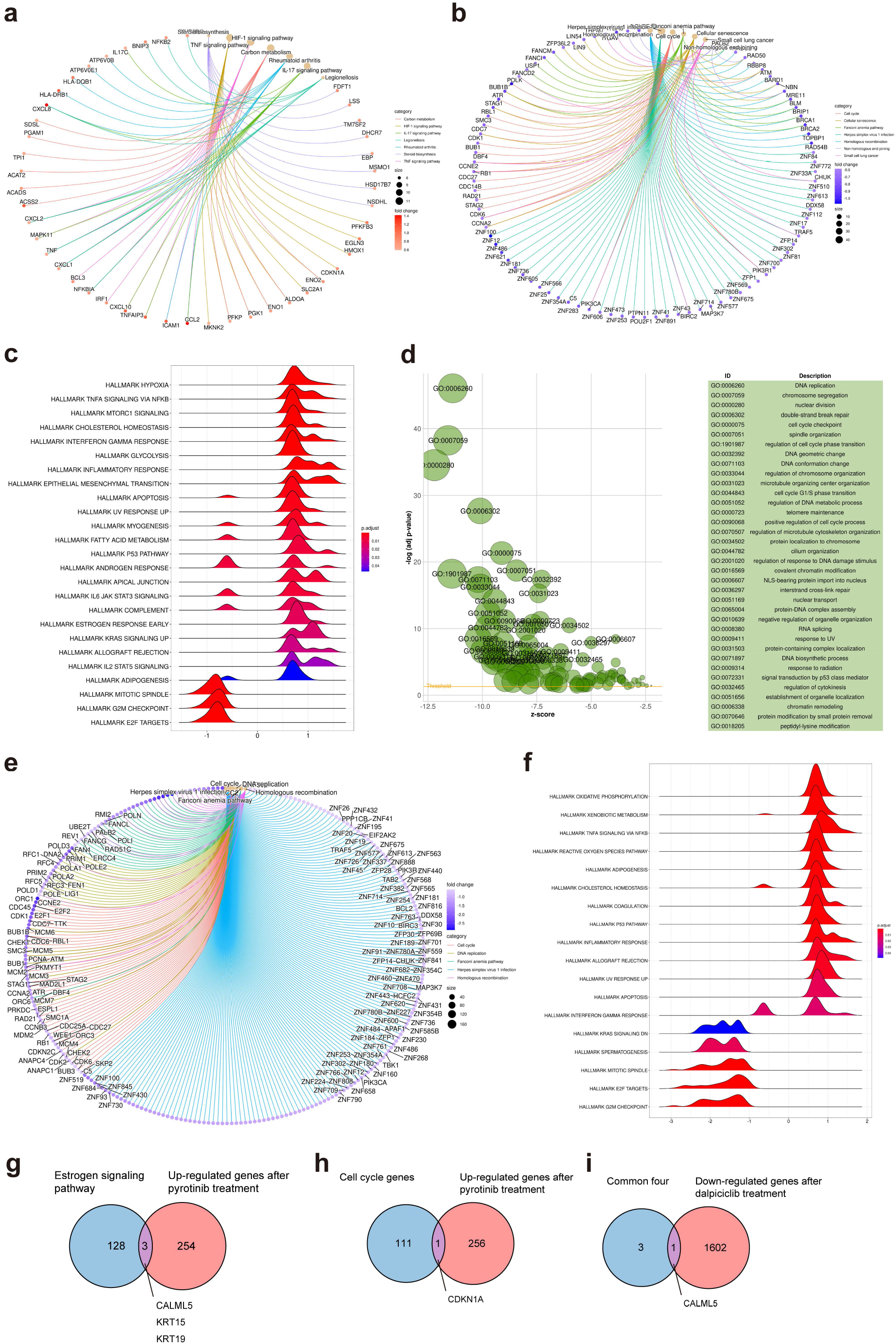
Bioinformatic analysis revealed dalpiciclib and pyrotinib blocking HER2 pathway and cell cycle in BT474 cells synergistically. a-b: Signaling pathway enrichment analysis of mRNA changes of BT474 cells treated with pyrotinib compared to BT474 cells treated with 0.1%DMSO. c: GSEA analysis of mRNA changes of BT474 cells treated with pyrotinib compared to BT474 cells treated with 0.1%DMSO. d-e: Signaling pathway enrichment analysis of mRNA changes of BT474 cells treated with pyrotinib+ tamoxifen+dalpiciclib compared to BT474 cells treated with pyrotinib+ tamoxifen. f: GSEA analysis of mRNA changes of BT474 cells treated with pyrotinib+ tamoxifen+ dalpiciclib compared to BT474 cells treated with pyrotinib+ tamoxifen. g: Intersection of genes which was upregulated after pyrotinib treatment and belonged to estrogen receptor signaling pathway (Genes belonged to estrogen receptor signaling pathway was provided in Figure 3 source data 1). h: Intersection of genes which was upregulated after pyrotinib treatment and belonged to cell cycle genes (Genes belonged to cell cycle gens were provided in Figure 3 source data 2). i: Intersection of the four genes which was upregulated after pyrotinib treatment and was downregulated after the introduction of dalpiciclib (genes which was upregulated after pyrotinib treatment and was downregulated after the introduction of dalpiciclib were provided in Figure 3 source data 3 and Figure 3 source data 4).

We then investigated the alteration of the gene expression profiles between breast tumor cells treated with triple-combined drugs (pyrotinib, tamoxifen, and dalpiciclib) and those treated with the dual-combined drugs (pyrotinib and tamoxifen) via gene enrichment analyses. The results suggested that the addition of dalpiciclib markedly reduced cell cycle progression. This was characterized by the enrichment of the cell cycle and the DNA replication process (Figure 3e). The GSEA results further indicated that the progression of the cell cycle was impeded by the enrichment of the gene sets, including MITOTIC SPINDLE and G2M CHECKPOINT (Figure 3f).

The activation of the ER pathway might be involved in the effect of pyrotinib on HER2^+^/HR^+^ breast cancer cells; therefore, intersection analyses were performed to confirm this. As shown in Figure 3g, *CALML5, KRT15,* and *KRT19* are the common genes shared between the two sets, the upregulated genes treated with pyrotinib compared to DMSO control group and the genes belonging to the estrogen signaling pathway. Since dalpiciclib is a cell cycle blocker, we also analyzed the common genes involved in the upregulation of the genes and the cell cycle progression after pyrotinib treatment. *CDKN1A* was the only shared gene in these two sets (Figure 3h). We then investigated whether any of the above-mentioned genes were upregulated with the use of pyrotinib and whether this could be abrogated with the introduction of dalpiciclib, which may serve as a potential risk factor in the treatment of HER2^+^HR^+^ breast cancer. The results showed that only one factor, *CALML5*, was the common gene (Figure 3i).

### CALML5 is a potential risk factor in the treatment of HER2^+^HR^+^ breast cancer

Western blot analyses and bioinformatic analyses were conducted to verify the changes in the signaling pathways. The western blot analyses showed that while the introduction of tamoxifen did not significantly affect the expression of HER2 and partially inhibited the HER2 downstream pathway (AKT-mTOR signaling pathway), it did not significantly affect the phosphorylation of Rb (Figure 4a). In contrast, the combination of pyrotinib and dalpiciclib showed similar inhibition of HER2 downstream pmTOR as the combination of pyrotinib and tamoxifen (Figure 4a). However, the combination of pyrotinib and dalpiciclib significantly reduced pRb expression and pCDK4(Thr172) expression (Figure 4a). In addition, cell arrest analyses of the different drug combinations were performed. As shown in Figure 4 b, compared with the cells treated with pyrotinib or tamoxifen, the introduction of dalpiciclib significantly increased the number of cells arrested in the G1/S phase. This confirmed the synergistic inhibition of cell proliferation by dalpiciclib and pyrotinib.

**Figure 4.**
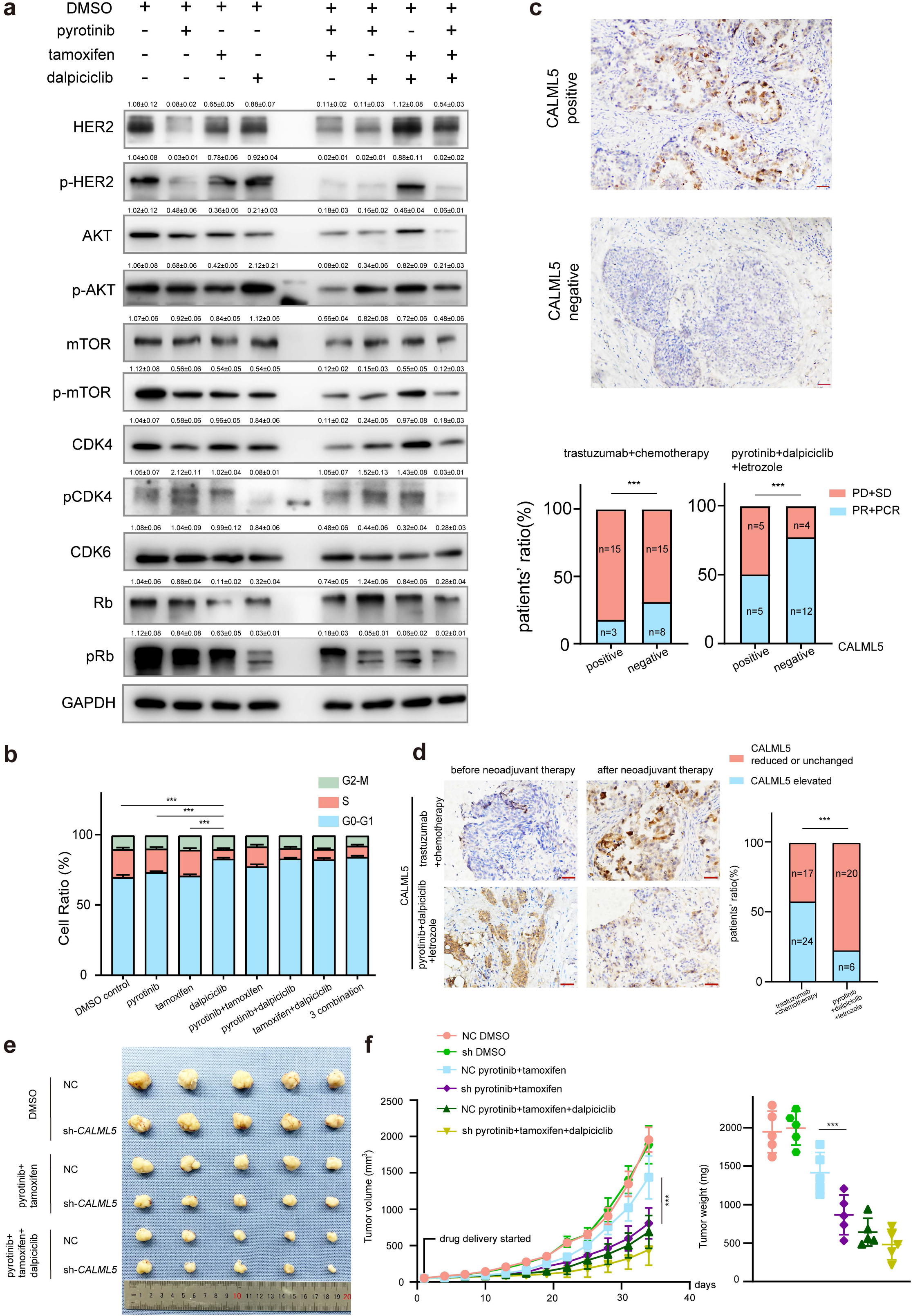
CALML5 could serve as a potential risk factor in the treatment of HER2^+^HR^+^ breast cancer. a: Western blot analysis of HER2 signaling pathway and cell cycle pathway in BT474 cells treated with different drugs or their combination. (This assay was performed in triplicates independently). b: Cell cycle analysis in BT474 cells treated with different drugs or their combination. (Data was presented as mean ± SDs, \*\*\**P*<0.001 using repeated Anova test; all the assays were performed independently in triplicates). c: Representative views of CALML5 positive/negative tissue. The difference of PR+PCR ratio and PD+SD ratio in patients who received anti-HER2 therapy (trastuzumab)+chemotherapy (docetaxel+carboplatin) or pyrotinib+dalpiciclib+l etrozole regarding on their expression of CALML5. (\*\*\**P*<0.001 using chi square test). d: Representative views of CALML5 positive/negative tissue. Ratio of patients with elevated or decreased CALML5 after receiving anti-HER2 therapy (trastuzumab)+chemotherapy (docetaxel+carboplatin) or pyrotinib+dalpiciclib+letrozole. (\*\*\**P*<0.001 using chi square test). e: Representative views of xenograft tumors derived from BT474 NC (NC stands for negative control) or BT474 sh cell lines treated with different drug combination. (\*\*\**P*<0.001 using Student’s t-test) f: Growth curves and tumor weight of xenograft tumors derived from BT474 NC or BT474 sh cell lines treated with different drug combination. (n=5 in each group, \*\*\**P*<0.001 using Student’s *t*-test) Raw gels were provided in Figure 4 source data 1, statistical data was provided in Figure 4 source data 2, original files of cell cycle analysis were provided in Figure 4 source data 3.

To verify whether CALML5 could be a potential risk factor of treatment responsiveness in clinical practice, clinical samples were collected from HER2^+^/HR^+^ patients before and after neoadjuvant therapy (anti-HER2 therapy(trastuzumab) + chemotherapy(docetaxel+carboplatin) or anti-HER2 therapy(pyrotinib) + CDK4/6 inhibitor(dalpiciclib)+endocrine therapy(letrozole))(Table 2). Immunohistochemical staining of CALML5 showed that the CALML5-positive cells indicated worse drug sensitivities and lower probabilities of achieving pathological complete response (pCR) and partial response (PR) in patients receiving neoadjuvant therapy (Figure 4c). However, pyrotinib + letrozole+dalpiciclib displayed better pCR and PR rates than trastuzumab + chemotherapy (docetaxel+carboplatin) in patients with CALML5-positive cells (Figure 4c). Moreover, the positive rate of CALML5 decreased after pyrotinib + letrozole+dalpiciclib treatment (Figure 4d), consistent with the results of the bioinformatic analyses. Furthermore, xenografts models derived from BT474 cells were also used to test the fuction of CALML5 in models using pyrotinib+tamoxifen or pyrotinib+tamoxifen+dalpiciclib. After knock down *CALML5* (Figure 4-figure supplement 1a), the tumor seemed to be more sensitive to the treatment of pyrotinib+tamoxifen (Figure 4 e and f) and it showed similar response compared to the group treated with 3 drug combination (Figure 4e and f). Hence, using clinical specimens as well as in vivo models, we found that the expression of CALML5 might be the potential risk factor in the treatment of HER2^+^HR^+^ breast cancer and the introduction of CDK 4/6 inhibitor could abrogate this.

## Discussion

Until now, the combination of antiHER2 therapy and chemotherapy have been the major treatment strategies for treatment of HER2^+^/HR^+^ breast cancer (*Gianni et al., 2012*; *Schneeweiss et al., 2013*). Although pCR and DFS improve with the use of the combination of anti-HER2 therapy and chemotherapy, the strong adverse effects of chemotherapy cannot be ignored (*Maguire et al., 2021*). Moreover, clinical data showed that the addition of anti-estrogen receptor drugs in the treatment regimen of HER2^+^/HR^+^ breast cancer did not provide additional advantages in the pCR rates and DFS (*Harbeck et al., 2017; Rimawi et al., 2017*). Hence, with the rapid development of small-molecule drugs such as tyrosine kinase inhibitors (TKIs) and CDK4/6 inhibitors, additional chemo-free strategies are being developed for the treatment of HER2^+^/HR^+^ breast cancer (*Gianni et al., 2018; Pascual et al., 2021; Saura et al., 2014*). In the recent MUKDEN 01 clinical trial (NCT04486911), the combination of pyrotinib, letrozole and dalpiciclib achieved satisfactory clinical response in HER2^+^HR^+^ patients with minimal adverse effects and offered novel chemo-free neoadjuvant therapy for HER2^+^HR^+^ patients(*Niu et al., 2022*). The molecular mechanism how the combination of pyrotinib, letrozole and dalpiciclib achieved optimal therapeutic effect remained further investigation.

In our study, we found that the combination of tamoxifen and pyrotinib was less effective in cytotoxicity than the combination of pyrotinib and dalpiciclib in BT474 cancer cells. This was anomalous since the two blocking agents of HER2 and ER were expected to inhibit their crosstalk and achieve better responses. To explore the potential mechanisms, we investigated the crosstalk between HER2 and the ER. After degrading HER2 with pyrotinib, ER was found to relocate to the cell nucleus, enhancing the function of ER which was consistent with the findings of Kumar et al and Yang et al (*Kumar et al., 2002*; *Yang et al., 2004*). We believe that the anti-HER2 mediated ER redistribution caused the enhanced ER function, leading to the relatively low cytotoxic efficacy of the combination of pyrotinib and tamoxifen in the treatment of HER2^+^/HR^+^ cells. Moreover, we found that the introduction of dalpiciclib to pyrotinib significantly decreased the total and nuclear expression of ER, partially abrogated the ER activation caused by pyrotinib. This may be the underlying mechanism by which the addition of dalpiciclib could achieve better response in the in vitro and in vivo studies.

Furthermore, using mRNA-seq and bioinformatics analyses, CALML5 was selected as a potential risk factor in the treatment of HER2^+^HR^+^ breast cancer. CALML5, known as calmodulin-like 5, is a skin-specific calcium-binding protein that is closely related to keratinocyte differentiation (*Mehul et al., 2001*). A previous study showed that the high expression of CALML5 was strongly associated with better survival in patients with head and neck squamous cell carcinomas (*Wirsing et al., 2021*). Misawa et al. (*Misawa et al., 2020*) reported that the methylation of CALML5, led to its downregulation, and this showed a correlation with HPV-associated oropharyngeal cancer. Moreover, the ubiquitination of CALML5 in the nucleus was found to play a role in the carcinogenesis of breast cancer in premenopausal women (*Debald et al., 2013*). Our results suggested that HER2^+^/HR^+^ breast cancer patients with positive CALML5 may be relatively drug resistant to anti-HER2 therapy (pyrotinib or trastuzumab) and the introduction of dalpiciclib might overcome this and offer better therapeutic effects. However, the underlying mechanism of CALML5 in breast cancer requires further investigation.

In conclusion, our study investigated the underlying synergistic mechanism for the combination of pyrotinib, letrozole and dalpiciclib in the MUKDEN 01 clinical trial (NCT04486911). We displayed the novel role of the dalpiciclib in HER2^+^/HR^+^ breast cancer, provided evidence that CALML5 may serve as a potential risk factor in the treatment of HER2^+^HR^+^ breast cancer and the introduction of dalpiciclib might overcome this.

## Materials and methods

### Clinical specimens

A total of 198 HR^+^/HER2^+^ patients who received neoadjuvant therapy were enrolled in this study to evaluate the status of ER and CALML5, of which 26 patients were from the clinical trial (NCT04486911, An open-label, multicentre phase II clinical study of pyrotinib maleate combined with CDK4/6 inhibitor and letrozole in neoadjuvant treatment of stage II-III triple positive breast cancer), 41 patients received anti-HER2 therapy(trastuzumab)+chemotherapy(docetaxel+carboplatin) and 131 patients only received chemotherapy(docetaxel+carboplatin). The sample size was calculated based on the four interrelated statistics in the Null Hypothesis Significant Test (NHST): sample size, effect size, alpha level, and statistical efficacy. The clinical information and specimens were analyzed to determine the impact of endocrine therapy on prognosis.

The study was approved by the Institutional Ethics Committee and complied with the principles of the Declaration of Helsinki and Good Clinical Practice guidelines of the National Medical Products Administration of China. Informed consent was obtained from all the participants.

### Cell lines and cell cultures

BT474 were purchased from the American Type Culture Collection (ATCC, Manassas, VA, USA). The human HER2^+^/HR^+^ breast cancer cell line BT474 was cultured in RPMI1640 culture medium supplemented with 10% fetal bovine serum (FBS).

### Chemicals and antibodies

Pyrotinib (SHR1258) and dalpiciclib (SHR6390) were kindly provided by Hengrui Medicine Co., Ltd. Tamoxifen (HY-13757A) and Trastuzumab was purchased from MCE company. Compounds were dissolved in dimethylsulfoxide (DMSO) at a concentration of 10 mM and stored at −20 °C for further use. Trastuzumab were dissolved and used according to manufacturer’s instructions. The following antibodies were purchased from Cell Signaling Technology (Beverly, MA, USA): ER, p-HER2 (Tyr 1221/1222), HER2, p-Akt (Ser473), AKT, p-mTOR, mTOR, pRb (Ser 780), Rb, CDK4, CDK6, Ubi, Lamin A, HSP90 and GAPDH. The pCDK4(Thr172) antibody was purchased from Absin Technologies (Shanghai, China).

### Cell viability assays and drug combination studies

CCK cell viability assays were (Cofitt life science) used to quantify the inhibitory effect of the different treatments. Cells were seeded in 96-well plates at a density of 5000 cells/well and treated the next day with DMSO, pyrotinib, trastuzumab, tamoxifen, dalpiciclib, or both drugs in combination for 48 h. The combination index (CI) values of different drugs were calculated using CompuSyn (ComboSyn Inc.). The CI values demonstrated synergistic (<1), additive (1–1.2), or antagonistic (>1.2) effects of the two-drug combinations. The drug sensitivity experiments were performed three times independently.

### Cell cycle analyses

The cells were starved in culture medium supplemented with 2% serum for 24 h before treatment. Treatments included DMSO (0.1%), pyrotinib (10 nM), dalpiciclib (8 μM), tamoxifen (5 μM), or different combinations of drugs. After treatment for 24 h, cells in different treating groups were trypsinized, washed with PBS, fixed in 70% ethanol, and incubated overnight at 4 °C. Next day, cells were collected, washed, and re-suspended in PBS at a concentration of 5 × 10^5^ cells/mL. The cell solutions were then incubated with a RNase and propidium iodide (PI) solution for 30 min at room temperature without exposure to light, and analyzed using a flow cytometer (BD FACS Calibur) according to the manufacturer’s instructions. This assay was performed in triplicates.

### Colony formation assays

Cells were seeded in 6-well plates at a density of 1000 cells/well. The cells were treated with DMSO (0.1%), pyrotinib (10 nM), tamoxifen (5 μM), dalpiciclib (8 μM), or a combination of the two or three agents. During the process, the culture medium was renewed every three days. After 14 days, the colonies were fixed and stained with crystal violet. Clusters of more than eight cells were counted as colonies. This assay was performed in triplicates independently.

### Western blot analysis

Cells were lysed using a cell lysis buffer (Beyotime, Shanghai, China). The total proteins were extracted in a lysis buffer (Beyotime, Shanghai, China), and the nuclear proteins were extracted using a nuclear protein extraction kit (Beyotime), in which protease inhibitor (HY-K0010; MCE) and phosphatase inhibitor (HY-K0021; MCE) were added. Protein concentrations were determined using a Pierce BCA Protein Assay Kit (Thermo Fisher Scientific, Waltham, MA, USA) according to the manufacturer’s instructions. The proteins from the cells and tissue lysates were separated using 10% SDS-PAGE and 6% SDS-PAGE, respectively, and then transferred to polyvinylidene fluoride (PVDF) membranes. The immunoreactive bands were detected using enhanced chemiluminescence (ECL). The western blot analysis was performed in triplicates independently.

### Co-Immunoprecipitation assay

BT474 cells treated with different drugs were lysed using a cell lysis buffer (Beyotime, Shanghai, China). in which protease inhibitor (HY-K0010; MCE) and phosphatase inhibitor (HY-K0021; MCE) were added. Protein concentrations were determined using a Pierce BCA Protein Assay Kit (Thermo Fisher Scientific, Waltham, MA, USA) according to the manufacturer’s instructions. Lysates were clarified by centrifugation, incubated with primary ER antibodies (#8644; Cell Signaling Technologies) overnight at 4°C, and incubated with protein A/G coupled sepharose beads (L1721; Santa Cruz Biotechnology) for 2 hours at 4°C. Bound complexes were washed 3 times with cell lysis buffer and eluted by boiling in SDS loading buffer. Bound proteins were detected on 6% SDS-PAGE followed by immunoblotting. The immunoreactive bands were detected using enhanced chemiluminescence (ECL).

### Immunofluorescence assays

The cellular localization of different proteins was detected using immunofluorescence. Briefly, the cells grown on glass coverslips were fixed in 4% paraformaldehyde at room temperature for 30 min. Cells were incubated with the respective primary antibodies for 1 h at room temperature, washed in PBS, and then incubated with 590-Alexa-(red) secondary antibodies (Molecular Probes, Eugene, OR, USA). We used 590-Alexa-phalloidin to localize the ER. The nuclei of the cells were stained with DAPI and color-coded in blue. The images were captured using an immunofluorescence microscope (Nikon Oplenic Lumicite 9000). The distribution ratio of ER was calculated manually by randomly chosen 5 views in 400magnification. The immunofluorescence assay was performed in triplicates independently.

### Immunohistochemical staining

The clinical samples were fixed in 4% formaldehyde, embedded in paraffin, and sectioned continuously at a thickness of 3 μm. The paraffin sections were deparaffinized with xylene and rehydrated using a graded ethanol series. They were then washed with tris-buffered saline (TBS). After these preparation procedures, the sections of each sample were incubated with the primary anti-ER antibody (Abcam Company, ab32063), anti-HER2 antibody (Abcam Company, ab134182), and anti-CALML5 antibody (Proteintech, 13059-1-AP) at 4 °C overnight. The next day, they were washed three times with TBS and incubated with a horseradish peroxidase (HRP)-conjugated secondary antibody (Gene Tech Co. Ltd., Shanghai, China) at 37 °C for 45 min, followed by immunohistochemical staining using a DAB kit (Gene Tech Co. Ltd.) for 5–10 min.

### Evaluation of the ER and HER2 status

The ER and HER2 statuses of patients who received neoadjuvant therapy were evaluated by a pathologist from a Shenjing affiliated hospital. The clinical specimens before and after the neoadjuvant therapy were evaluated. The analyses of the elevation or decline in ER statuses were based on these pathological reports. The 2+ of HER2 was detected by immunohistochemistry as well as a FISH test positive report.

### mRNA-seq and differential gene expression analysis

BT474 cells were treated with 1%DMSO, pyrotinib (10 nM), tamoxifen (5 μM), dalpiciclib (8 μM), pyrotinib+tamoxifen, pyrotinib+dalpiciclib, tamoxifen+dalpiciclib and combination of 3 drugs, respectively. Each group was performed in triplicate an treated with drugs for 48 hours. After the treatment, the mRNAs in these cells were extracted using RNAiso Plus (Takara, Cat:9109) and then sequenced by Biomarker Techonologies using Illumina sequencing technology. The differential gene expression analysis was performed using online tools (http://www.biomarker.com.cn/biocloud), differential expressed genes were defined as Log2 Foldchange>0.5, P value <0.05. As for the gene set of estrogen signaling pathway and cell cycle genes, genes sets were downloaded from KEGG database.

### Gene enrichment analysis

Gene annotation data in the GO and KEGG databases and R language were used for the enrichment analysis. Only enrichment with q-values less than 0.05 were considered significant.

### GSEA

The hallmark gene sets in the Molecular Signatures Database were used for performing the GSEA; only gene sets with q-values less than 0.05 were considered significantly enriched.

### Stably knock down of CALML5 in BT474 cell line

The sh-CALML5 lentivirus was synthesized by Genechem Technologies. BT474 cells were cultured in a 6-well plate and transduced with shRNAs targeting human CALML5 or NC (negative control). The sequences for sh - *CALML5* were 5’-ACGAGGAGTTCGCGAGGAT-3’ (sequence 1),5’-AAATCAGCTTCCAGGAGTT-3’(sequence 2) and 5’-GAAACTCATCTCCGAGGTT-3’(sequence 3). The sequence for sh-NC was 5’-GCAGTGAAAGATGTAGCCAAA-3’.

### Animal studies

Four-to five-week-old female NOD scid mice were maintained in the animal husbandry facility of a specific pathogen free (SPF) laboratory. All experiments were performed in accordance with the Regulations for the Administration of Affairs Concerning Experimental Animals and were approved by the Experimental Animal Ethics Committee of the China Medical University.

Subcutaneous injections of 1×10^7^ BT474 NC cells or BT474 sh-*CALML5* cells were performed to induce tumors. 2 weeks after tumor cell inoculation, tumor volume was measured every 3 days and calculated as V = 1/2 (width^2^ × length).

As for drug sensitivity test, pyrotinib, tamoxifen and dalpiciclib was administrated when after 2 weeks of tumor inoculation. Mice inoculated with BT474 NC or BT474 sh-*CALML5* cells were randomly assigned to one of 3 groups (n=5 each, total number=30). Mice carried xenograft tumors were treated by intraperitoneal injection for 28 days with vehicle (1% DMSO dissolved in normal saline/2d), pyrotinib (20mg/kg every 3 day), tamoxifen (25mg/kg every 3day) and dalpiciclib (75mg/kg every half a week). When the drug was continuously delivered for 32 days, mice were humanely euthanized and tumors were dissected and analyzed.

### Statistical analysis

All the descriptive statistics were presented as the means ± standard deviations (SDs). The differences between two groups were analyzed by Student’s t tests and the differences among groups were analyzed by repeated Anova tests. The differences between percentage data were analyzed using chi square test. Kaplan-Meier methods were used to compute the survival analysis and *P*-value was obtained by log-rank test. The statistical analyses were performed using IBM SPSS version 22 (SPSS, Armonk, NY, USA) and GraphPad Prism version 7. The statistical significance of the differences between the test and control samples was assessed at significance thresholds of \**P* < 0.05, \*\**P* < 0.01 and \*\*\**P* < 0.001.

## Authors Contributions

J.B., Y.Z., L.S., X.Q., Y.W., X.J., D.W., H.L., and Q.M. conceptualized the study, performed the experiments, and analyzed the data. B.K. performed the bioinformatic analysis. Y.Z. and N.N. provided the clinical data and samples. C.L. designed the entire study and wrote the manuscript.

## Conflict of interest

The authors declare no conflicts of interests. H.L is affiliated with Jiangsu Hengrui Pharmaceuticals Co. Ltd and the author has no other competing interests to declare.

## Funding

This study was supported by the National Natural Science Foundation of China (#U20A20381, #81872159)

## Supplementary

**Figure 1-figure supplement 1.**
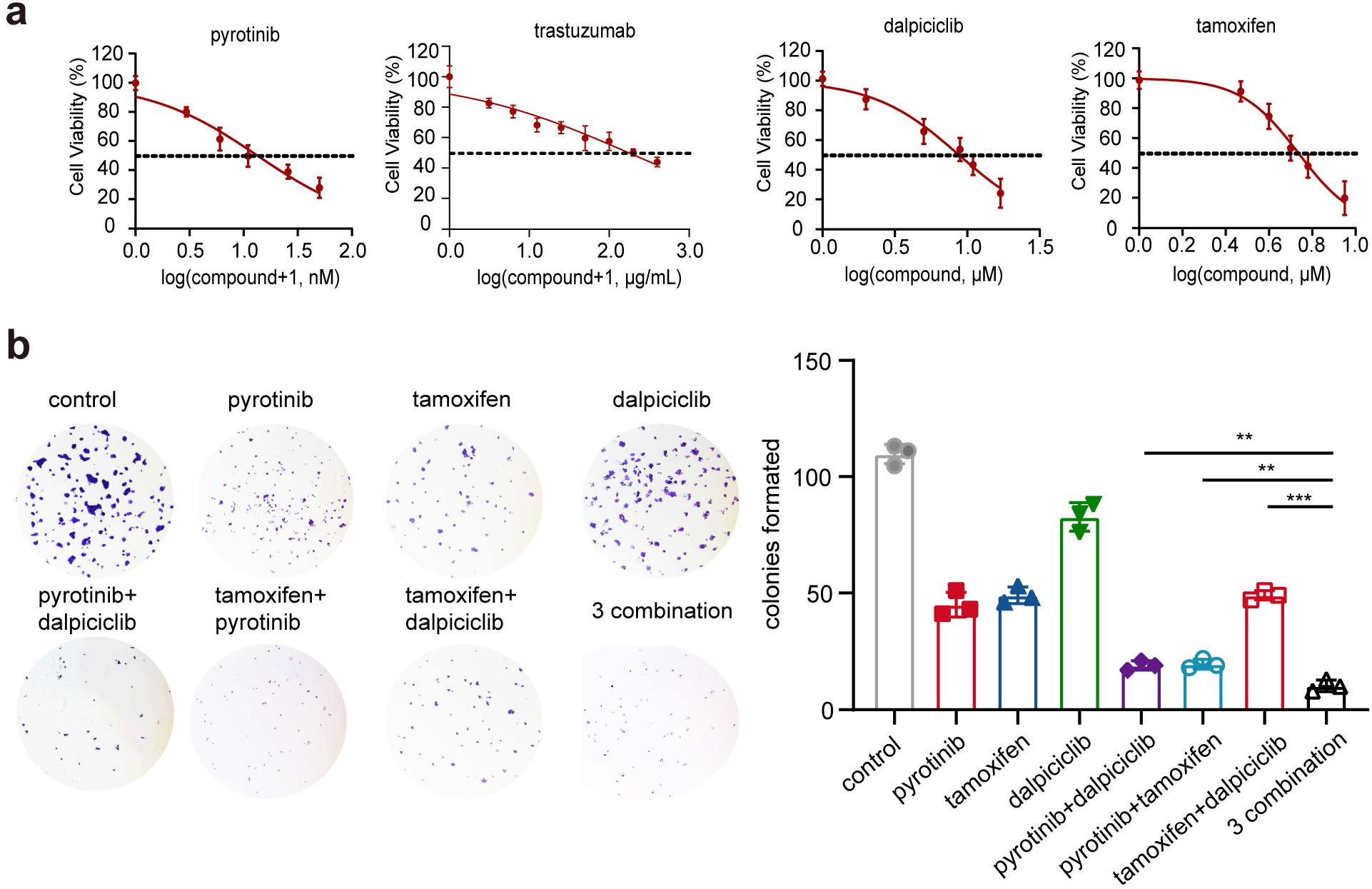
a: Drug sensitivity analysis of pyrotinib, tamoxifen and dalpiciclib in BT474 cells. (Data was presented as mean ± SDs, all the assays were performed independently in triplicates). b: Colony formation assay of BT474 cells treated with different drugs. (Data was presented as mean ± SDs, \*\**P*<0.01 and \*\*\**P*<0.001 using repeated Anova test; all the assays were performed independently in triplicates) Statistical data was provided in Figure 1-figure supplement 1 source data 1.

**Figure 2-figure supplement 1.**
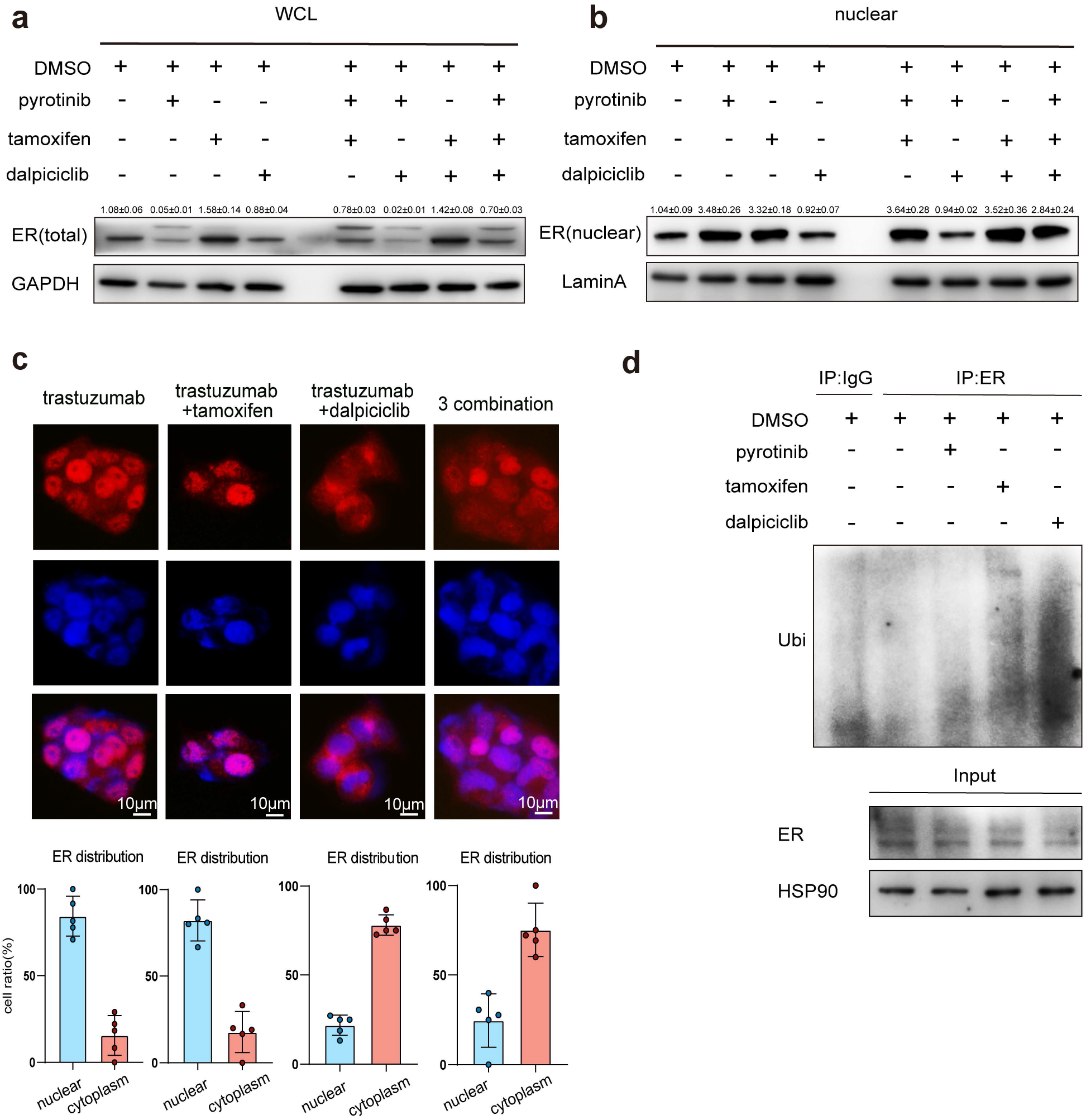
a-b: Total ER expression and nuclear ER expression in BT474 cells treated with different drugs. (This assay was performed in triplicates independently). c: Distribution of estrogen receptor in BT474 cell line after different drug (trastuzumab, tamoxifen and dalpiciclib) treatment. (The distribution ratio of ER was calculated manually by randomly chosen 5 views in 400 magnification. All the assays were performed independently in triplicates, Figure 2-figure supplement 1 source data 2). d: The ubiquitination of ER in BT474 cells after the treatment of DMSO, pyrotinib, tamoxifen and dalpiciclib. Raw gels were provided in Figure 2-figure supplement 1 source data 1. Statistical data was provided in Figure 2-figure supplement 1 source data 2.

**Figure 4-figure supplement 1.**
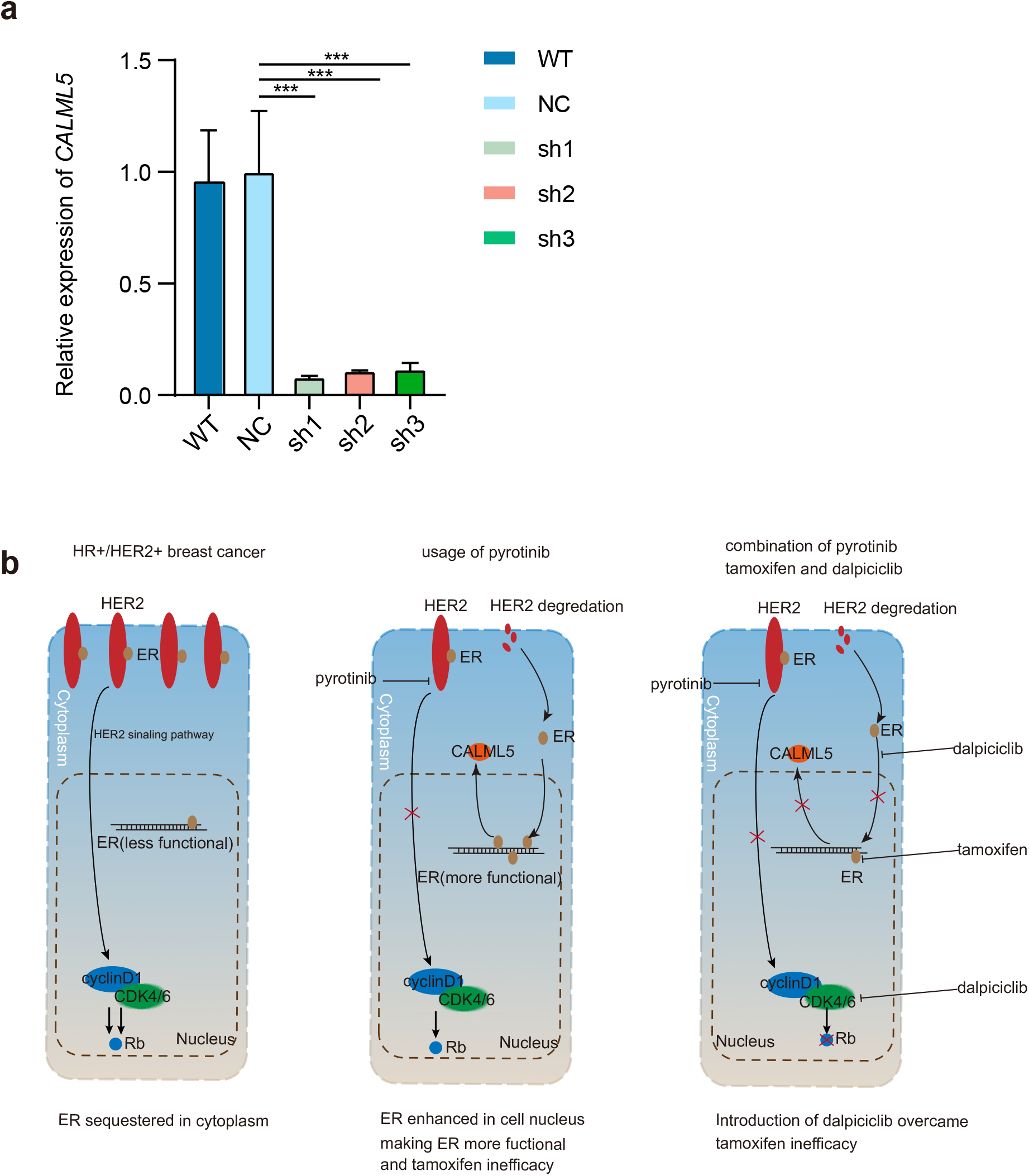
a: The efficacy of the sh-*CALML5* lentivirus detected by qRT-PCR and the sh1 sequence was used in the xenograft study, NC stands for negative control. (Data was presented as mean ± SDs, \*\*\**P*<0.001 using repeated Anova test; all the assays were performed independently in triplicates) Statistical data was provided in Figure 4-figure supplement 1 source data 1. b: The introduction of dalpiciclib to pyrotinib could significantly decrease the total and nuclear expression of ER, thus partially abrogate the ER activation caused by pyrotinib and CALML5 could be served as a potential marker of ER activation after the treatment of pyrotinib.

## Source data

Figure 1 source data 1

Statistical data of Figure 1

Figure 2 source data 1

Statistical data of Figure 2

Figure 3 source data 1

Gene list in ER signaling pathway summarized by KEGG database for Figure 3 g.

Figure 3 source data 2

Gene list in cell cycle genes summarized by KEGG database for Figure 3 h.

Figure 3 source data 3

Up-regulated genes after pyrotinib treatment compared to DMSO treatment for Figure 3 g and i.

Figure 3 source data 4

Down-regulated genes after dalpiciclib treatment compared to DMSO treatment for Figure 3 h and i.

Figure 4 source data 1

Original files for the gels in Figure 4 a.

Figure 4 source data 2

Histograms of the cell cycle analysis in Figure 4 b.

Figure 4 source data 3

Statistical data for Figure 4.

Figure 1-figure supplement 1 source data 1

Statistical data for Figure 1-figure supplement 1.

Figure 2-figure supplement 1 source data 1

Original gels for Figure 2-figure supplement 1 a, b and d.

Figure 2-figure supplement 1 source data 2

Statistical data for Figure 2-figure supplement 1.

Figure 4-figure supplement 1 source data 1

Statistical data for Figure 4-figure supplement 1.

## Notes

### Competing Interest Statement

The authors have declared no competing interest.

